# Temperature effects on interspecific eavesdropping in the wild

**DOI:** 10.1101/2024.08.28.610172

**Authors:** Sarina M. Rossi, Kasey D. Fowler-Finn, David A. Gray

## Abstract

Mating signals are targets of conspecific signal recognition and sexual selection, but are also subject to abiotic temperature effects and to biotic interspecific eavesdroppers. In crickets, the male calling song becomes faster at warmer temperatures, and female crickets’ recognition of male song tracks temperature in a coordinated manner, termed ‘temperature coupling.’ But female crickets are not the only ecologically relevant listeners: some cricket species are parasitized by *Ormia ochracea*, a parasitoid fly which finds its cricket hosts by eavesdropping on male cricket song. How temperature affects parasitoid fly phonotaxis to song is largely unexplored, with only one previous study conducted under field conditions. Here we explore six possible patterns of thermal effects on fly responses to cricket song, including temperature coupling, using field playbacks of synthetic *Gryllus lineaticeps* songs designed to be species-typical at various temperatures. We find that temperature does affect fly response, but that the temperature deviation of songs from ambient does not impact numbers of flies caught. We extend this finding by comparing the temperatures of the air and ground to show that temperature coupling is unlikely to be effective given microhabitat variation and differential rates of cooling in the evening hours when flies are most active. Our results can be interpreted more broadly to suggest (i) temperature effects on intraspecific communication systems may be more tightly coupled than are effects on interspecific eavesdropping, and (ii) variation in thermal microhabitats in the field make it difficult to translate laboratory physiological responses to natural selection in the wild.

**Lay Summary:** Mating signals and signal recognition change with temperature, and sometimes mating signals are intercepted by predators or parasites. By using playbacks of cricket song in the wild, we show that temperature changes do affect the response of a parasitoid fly to cricket song. However, parasitoid responses are not tightly coupled to temperature induced changes in cricket song, in part due to unpredictable variation in microhabitat temperatures typical of crickets and flies.

## INTRODUCTION

Temperature effects on ectotherms are ubiquitous (Abram et al., 2017; González-Tokman et al., 2020), including effects on communication signals (Brandt et al., 2018; Conrad et al., 2017; Macchiano et al., 2019; Shimizu and Barth, 1996; Symes et al., 2017). As signals have both senders and receivers, temperature changes could disrupt communication if changes in temperature affected each differentially (Leith et al., 2020). However, if temperature induces parallel changes in both senders and receivers, then effective communication could be maintained across a range of ambient temperatures. Such parallel changes are generally referred to as ‘temperature coupling’ (Jocson et al., 2019). In the case of intraspecific communication, such as with mating signals, senders and receivers share all or most of their genetic architecture, so pleiotropy could in principle link temperature effects in a coordinated manner. The potential for conspecific temperature coupling has been studied extensively (Brenowitz et al., 1985; Doherty and Hoy, 1985; Gerhardt, 1978; Greenfield and Medlock, 2007; Jocson et al., 2019; Ritchie, 1992), mostly in the absence of direct evidence of pleiotropic effects on senders and receivers (Butlin and Ritchie, 1989), but see (Blankers et al., 2019; Shaw and Lesnick, 2009; Xu and Shaw, 2019, 2021). Alternatively, conserved properties of neurophysiology could produce parallel changes in senders and receivers (Greenfield and Medlock, 2007), even despite distantly shared ancestry. Such a circumstance could apply to ‘eavesdroppers’ which are typically heterospecific predators or parasites which use the signals of other species to target them for exploitation (Bernal and Page, 2023; Virant-Doberlet et al., 2019). Interspecific temperature coupling has been much less intensively studied than intraspecific coupling, but is supported by an experimental field study (Walker, 1993) and by recent laboratory work (Jirik et al., 2023), both conducted with an eavesdropping parasitoid fly *Ormia ochracea* which finds its cricket hosts using cricket mating signals as directional cues (Cade, 1975; Gray et al., 2019; Mason, 2021; Tinghitella et al., 2021; Vincent and Bertram, 2010).

Both Walker (1993) and Jirik et al. (2023) studied responses of *O. ochracea* from Florida, USA to pulse rate variation typical of temperature effects on the dominant host species of cricket in Florida, *Gryllus rubens*. Both studies found peak response of flies to pulse rates typical of *G. rubens* calls across a range of temperatures [ambient ± 4 ℃ in Walker (1993), and at 21, 25, or 29 ℃ in Jirik et al. (2023)]. In this study, we extend those works in several important ways: (i) we test responses of southern California *O. ochracea* to temperature dependent songs of the locally dominant host species of cricket, *G. lineaticeps*, (ii) we vary all parameters of cricket song that change with temperature (see Methods below), not just pulse rate, (iii) we consider 6 models of possible receiver response patterns (see below), not just temperature coupling, and (iv) we assess the likelihood of effective temperature coupling via comparison of air temperatures (experienced by flies) and ground temperatures (experienced by crickets).

### Possible patterns of receiver response

Although temperature coupling has been the most widely explored, it is not the only possible pattern of receiver response to temperature-induced variation in signals. We explore here six possible response patterns, but note that these six are not exhaustive. (i) Indifference: if the critical recognition feature of the signal is unaffected by temperature, then temperature induced variation in other features may be irrelevant. For example, dominant frequency of field cricket song is minimally affected by temperature (Martin et al., 2000), so if flies are highly responsive to any temporal pattern of ∼5 kHz pulses, across the fly’s range of active temperatures, then we would expect a random response pattern (Fig. 1a). (ii) Template: if receivers have a specific fixed template which responds best to a particular fixed stimulus, then that template would be revealed by the matching of an increase in ambient temperature with a negative deviation in stimulus relative to ambient. For example, a 75 pulse/s template would be equally stimulated by *G. lineaticeps* typical song at 25 ℃, or by a +4 ℃ song at 21 ℃, a +2 ℃ song at 23 ℃, a −2 ℃ song at 27 ℃, and a −4 ℃ song at 29 ℃ (Fig. 1b). (iii) Coupling: if receivers’ responses are matched to ambient temperature, and centered on the typical signal for that temperature, then peak response would be to the temperature-typical song across a range of ambient temperatures (Fig. 1c). (iv) Power: receivers may uniformly prefer greater stimulus levels typical of faster signalers regardless of ambient temperature (Fig. 1d). For example, both female *G. lineaticeps* crickets and *O. ochracea* flies are known to prefer faster chirp rates (Hennig et al., 2016; Wagner, 1996) which are associated with male crickets in better nutritional condition (Wagner and Hoback, 1999), but might also be males with a higher body temperature (Erregger et al., 2018). (v) Goldilocks: if receivers prefer intermediate ambient temperatures neither too hot nor too cold, and signals neither too fast nor too slow, then we would expect a ‘bullseye’ response pattern (Fig. 1e). (vi) Grandma: if receivers are indifferent to signal variation, but have a narrow range of preferred temperatures, then we would expect a response pattern centered just on that preferred ambient temperature (Fig. 1f). These idealized response scenarios can in principle be distinguished by different combinations of statistical model parameters (Table 1), however interpretation also requires examination of the parameter estimates (e.g., coupling described by a positive linear effect and a negative quadratic effect for playback deviation, and after ensuring that playback deviations are linearly transformed to positive values such that the squared quadratic effect is meaningful).

**Figure 1a-f.**
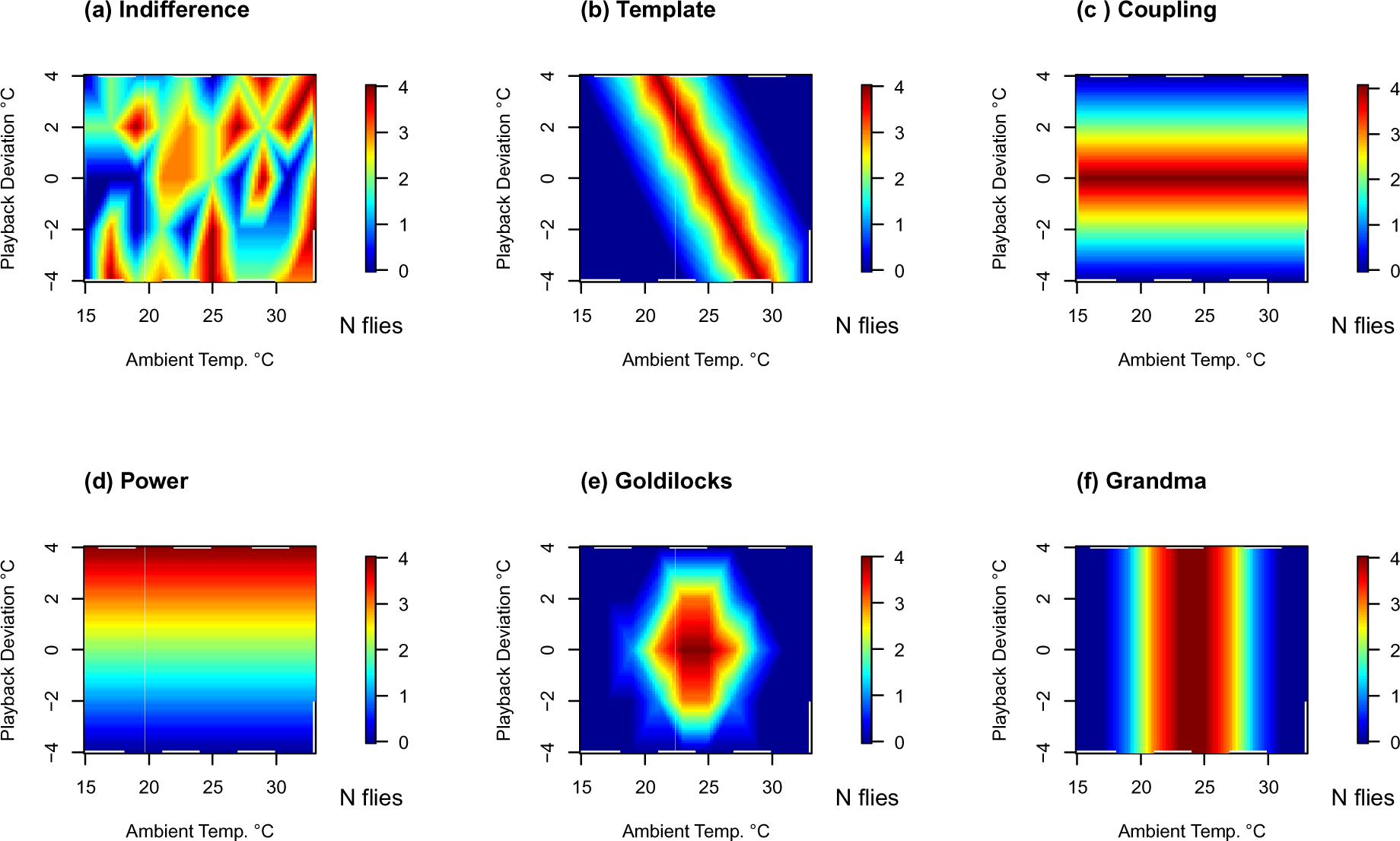
Hypothetical response patterns across a range of biologically plausible temperatures to songs which either match ambient temperature or deviate from it by ± 4 ℃.

**Table 1.** Statistical predictions useful to help distinguish among the six competing receiver response patterns (see text and Fig. 1).

|  | Intercept | Ambient<br>Temp. | Ambient<br>Temp.<sup>2</sup> | Playback<br>Deviation | Playback<br>Deviation<sup>2</sup> | Ambient<br>Temp. x<br>Playback<br>Deviation |
| --- | --- | --- | --- | --- | --- | --- |
| Indifference | * | NS | NS | NS | NS | NS |
| Template | * | * | * | NS | NS | * |
| Coupling | * | NS | NS | * | * | NS |
| Power | * | NS | NS | * | NS | NS |
| Goldilocks | * | * | * | * | * | * |
| Grandma | * | * | * | NS | NS | NS |

\* = predicted significant effect; NS = predicted not significant effect.

To test these ideas, we used field playbacks of temperature-matched cricket songs across naturally varying temperatures in the wild. Because organisms within the same habitat could experience very different thermal microhabitats (Leith et al., 2024; Scheffers et al., 2017), we also measured and compared air and ground temperatures to determine the feasibility of accurate parasitoid-host temperature coupling. Such data are rather scarce in the parasitoid-host literature (Flores-Mejia et al., 2016; Foray et al., 2011; Wang et al., 2012).

## METHODS

### Field sites

Fieldwork was conducted in August, September and October 2020 and 2021 in two semi-natural open spaces in the Santa Monica Mountains National Recreation Area in Los Angeles County, California, USA. We refer to these sites as Las Virgenes (LV, 34.104, −118.711), and Cheesebro (CH, 34.150, −118.735). Both sites are characterized by abandoned agricultural fields, now consisting primarily of non-native wild oat *Avena fatua* and black mustard *Brassica nigra* with native coast live oak *Quercus agrifolia* interspersed. Both *G. lineaticeps* and *O. ochracea* are abundant in the area (Beckers and Wagner, 2018; Dobbs et al., 2020; Paur and Gray, 2011a), and *G. lineaticeps* are the dominant local host for the fly (Gray et al., 2007). Research was conducted under US National Park Service permit SAMO-2020-SCI-0016.

### Playback exemplars

Playback exemplars were modified from *Gryllus lineaticeps* synthetic calls used by Gray et al. (2007), and were designed to represent species-typical calls at temperatures from 11 to 39 ℃ based on Maskell (1975) and DA Gray unpublished data (see Table 2). Songs were constructed from synthetic pulses created in CoolEdit 2000 (Syntrillium software) with a frequency sweep from 5000 to 4800 Hz with 800 Hz modulation at 5000 Hz modulation frequency. We broadcast the songs as 44.1 kHz 16 bit encoded .wav files from Wiwoo Sport Clip F3 Music Players connected via Bluetooth to Altec Lansing Mini H2O IP67 speakers.

**Table 2.** Parameters of *Gryllus lineaticeps* synthetic song exemplars with species-typical values for temperatures ranging from 11 to 39 ℃. [pd = pulse duration; ipi = inter-pulse interval; ppc = pulses/chirp; ici = inter-chirp interval; prate = pulse rate; cd = chirp duration; crate = chirp rate].

| TEMP | pd (ms) | ipi (ms) | ppc | ici (ms) | prate (p/s) | cd (ms) | crate (c/s) |
| --- | --- | --- | --- | --- | --- | --- | --- |
| 11 | 11 | 7.5 | 8 | 260 | 54 | 140.5 | 2.5 |
| 13 | 10.75 | 7 | 8 | 255 | 56 | 135 | 2.6 |
| 15 | 10.5 | 6.5 | 8 | 250 | 59 | 129.5 | 2.6 |
| 17 | 10.25 | 6 | 8 | 245 | 62 | 124 | 2.7 |
| 19 | 10 | 5.5 | 8 | 240 | 65 | 118.5 | 2.8 |
| 21 | 9.75 | 5 | 8 | 235 | 68 | 113 | 2.9 |
| 23 | 9.5 | 4.5 | 8 | 230 | 71 | 107.5 | 3.0 |
| 25 | 9.25 | 4 | 8 | 225 | 75 | 102 | 3.1 |
| 27 | 9 | 3.5 | 8 | 220 | 80 | 96.5 | 3.2 |
| 29 | 8.75 | 3 | 8 | 215 | 85 | 91 | 3.3 |
| 31 | 8.5 | 2.5 | 8 | 210 | 91 | 85.5 | 3.4 |
| 33 | 8.25 | 2 | 8 | 205 | 98 | 80 | 3.5 |
| 35 | 8 | 1.5 | 8 | 200 | 105 | 74.5 | 3.6 |
| 37 | 7.75 | 1 | 8 | 195 | 114 | 69 | 3.8 |
| 39 | 7.5 | 0.5 | 8 | 190 | 125 | 63.5 | 3.9 |

### Playback Trials

In any given trial, we simultaneously played five different synthetic *Gryllus lineaticeps* calls from five speakers placed one each underneath five slit traps designed for capturing *Ormia* (Walker, 1989). The five traps were arranged in a pentagon with 5 m sides; this approximates natural cricket densities in these populations (mean ± SD nearest neighbor distance of crickets at the LV site was 3.4 ± 2.7 m, DAG unpublished data, 2001). At the start of each trial, ambient air temperature was measured at 0.3 m above ground level in order to assign the appropriate call playbacks. Each trap was haphazardly assigned one of the following temperature-based songs: −4 ℃, −2 ℃, ambient, +2 ℃, +4 ℃. For example, if ambient at the start of the trial was 21 ℃, then the 5 playbacks were the species-typical songs for 17, 19, 21, 23, and 25 ℃. Call playbacks were broadcast simultaneously at 85 dB (re. 20 µPa) at 1 meter, measured with a RadioShack 33-2055 sound level meter (fast C-weighting) and a 5 kHz calibration tone that matched the peak amplitude of the call playbacks; 85 dB at 1 m is on the upper end of natural variation in most field crickets (Nandi and Balakrishnan, 2013; Simmons, 1988), and was used to increase the area over which flies could be attracted [see, e.g. Wagner (1996) and Dobbs et al. (2020) which, respectively, used 100 and 90 dB at 1 m for field fly phonotaxis playbacks for similar reasons]. On a few occasions, one of the speakers unexpectedly lowered in volume during a trial; on those occasions we briefly stopped the trial and re-calibrated the speakers.

Trials began shortly after sunset during the peak activity time of the fly (Beckers and Wagner, 2012; Cade et al., 1996). Each trial replicate lasted 15 minutes, and ambient temperature was measured again at the end of each trial. Traps were continuously checked, and any flies that landed on or entered the trap were captured and counted. We conducted from 1-3 trials on any given night, re-measuring ambient temperature and haphazardly re-assigning playbacks to speaker positions each trial. Captured flies were released at the end of each night after all trials for that night had concluded. We took this approach to avoid possible pseudo-replication from recapture of the same flies within one night; recapture of flies across different nights is unlikely based on a previous mark-recapture study with this same population of flies (Paur and Gray, 2011b).

### Correspondence between air and ground temperatures

In 2021, between trials on 6 different nights, we recorded the ground temperature of a total of 33 sites with characteristics commonly used by calling male crickets (e.g., soil cracks, rock edges) as well as the air temperature at ∼2m directly above each site using a Labcraft Digital Thermometer Temperature Probe with accuracy ± 0.1 ℃.

### Data analysis

Treatment of variables: The goal was to model the number of flies caught at each song as a function of (i) ambient temperature (hereafter ‘Ambient Temp.’, both linear and quadratic terms), (ii) deviation of the song model from ambient temperature (hereafter ‘Playback Deviation’, both linear and quadratic terms), and (iii) the Ambient Temp. x Playback Deviation interaction, all done after correcting for the covariates Site, Year, and Trial number within a single night. Site (LV and CH) and Year (2020 and 2021) are self-explanatory; Trial number (1st, 2nd, or 3rd within a night) was included because we temporarily collected all flies caught during each trial, and then released them all only at the conclusion of all trials at the end of each night. Therefore, it was reasonable to expect that the second or third trials on a given night might catch fewer flies than had the first trial that night. Ambient Temp. was defined as the average of the starting and ending temperatures for each trial. These were typically within only a degree or two of each other (mean ± SD drop in temperature during the 15 minute trials was 0.98 ± 0.89 ℃, N = 41 trials). Playback Deviation was then recalculated relative to that actual mean ambient temperature (i.e., the planned −4, −2, 0, +2, +4 ℃ deviations were adjusted to be the actual differences from the observed mean ambient temperature for that trial; this adjustment also corrects for our playback exemplars being spaced every 2 ℃). To facilitate the fit of quadratic terms, the Playback Deviations were then linearly transformed by +6 so that all values were positive, enabling meaningful use of the quadratic term.

Model selection: Given that the response variable was counts of flies, both Poisson and negative binomial models were considered and compared via likelihood ratio test using R version 4.3.1, packages *MASS* and *lmtest*. The negative binomial proved better (Chi-square 212.56, df = 1, p < 2.2e-16) and was a good fit to the data (residual deviance/df = 1.075).

## RESULTS

### Fly phonotaxis as a function of temperature

We conducted 41 total trials (1-3 per night, mean 1.56 ± 0.59 trials per night) over 20 nights (9 in 2020, 11 in 2021). The mean ± SD ambient temperature was 22.7 ± 4.7 °C [min 14.95, max 35.2]. In total, we caught 523 flies. After correcting for Site, Year, and Trial, we found that ambient temperature had a significant effect on the number of flies captured, but that playback song deviation from ambient did not affect fly response (Table 3). The temperature effect included significant positive linear and negative quadratic effects, such that fly response increased from low to mid temperatures, but then declined (Fig. 2). These results are consistent with the Grandma scenario: indifferent to song variation but with reduced responses at cold and hot temperatures.

**Figure 2:**
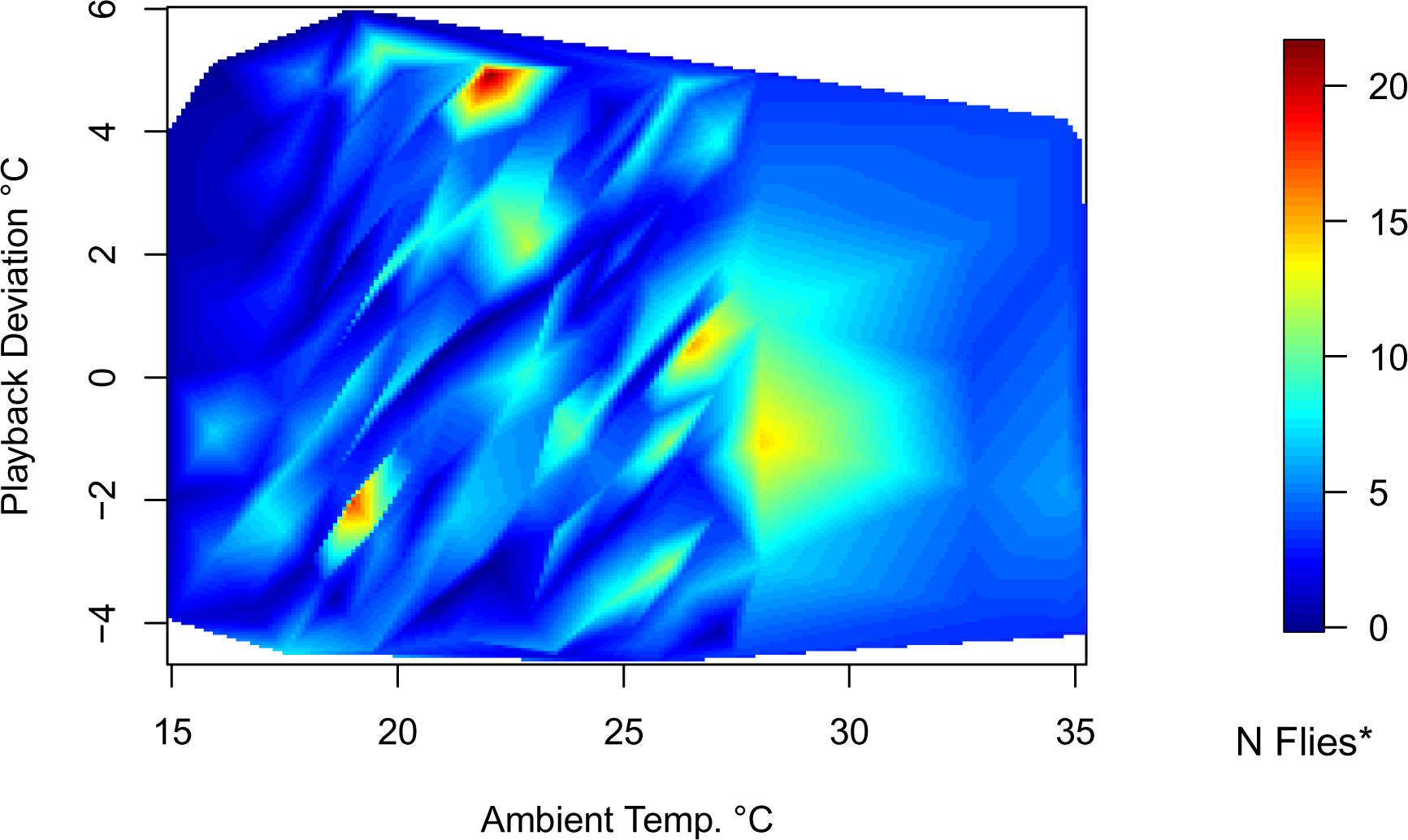
Numbers of flies caught as a function of ambient temperature and the deviation of song playback in ℃ from the typical song for a given ambient temperature. Note that fewer flies were caught at low and high temperatures, and that the numbers of flies caught appears random within a range of about 19 – 30 ℃, consistent with the ‘Grandma’ scenario. *For plotting, the number of flies was adjusted for covariates Site, Year, and Trial.

**Table 3.** Results of a General Linear Model with negative binomial error. To control for the covariates Site, Year, and Trial, the terms were entered in the order listed and Type.

| | Estimate ± SE | Z | Pr(>\|z\|) |
| --- | --- | --- | --- |
| Intercept | -2073 ± 390 | -5.316 | 1.06e-07 **** |
| Site | -1.053 ± 0.240 | -4.387 | 1.15e-05 **** |
| Year | 1.021 ± 0.193 | 5.291 | 1.22e-07 **** |
| Trial | -0.449 ± 0.167 | -2.697 | 0.00701 ** |
| Ambient Temp. | 0.883 ± 0.185 | 4.768 | 1.86e-06 **** |
| Ambient Temp.<sup>2</sup> | -0.017 ± 0.004 | -4.753 | 2.00e-06 **** |
| Playback Deviation | -0.031 ± 0.242 | -0.130 | 0.89675 NS |
| Playback Deviation<sup>2</sup> | -0.018 ± 0.012 | -1.495 | 0.13493 NS |
| Ambient Temp. x Playback Deviation | 0.009 ± 0.008 | 1.241 | 0.21454 NS |

NS = p > 0.05; \*\* = p < 0.01; \*\*\*\* = p < 0.0001

### Correspondence between air and ground temperatures

Air temperature as experienced by flies was a poor predictor of ground temperature at appropriate cricket calling sites (Fig. 3). Although air and ground temperatures were significantly and positively associated (N = 33, F_1,31_ = 6.28, P = 0.0176), the variation explained was low (R^2^ = 0.17), and the slope was significantly less than one (slope ± s.e. = 0.53 ± 0.21; versus slope = 1, P = 0.036).

**Figure 3.**
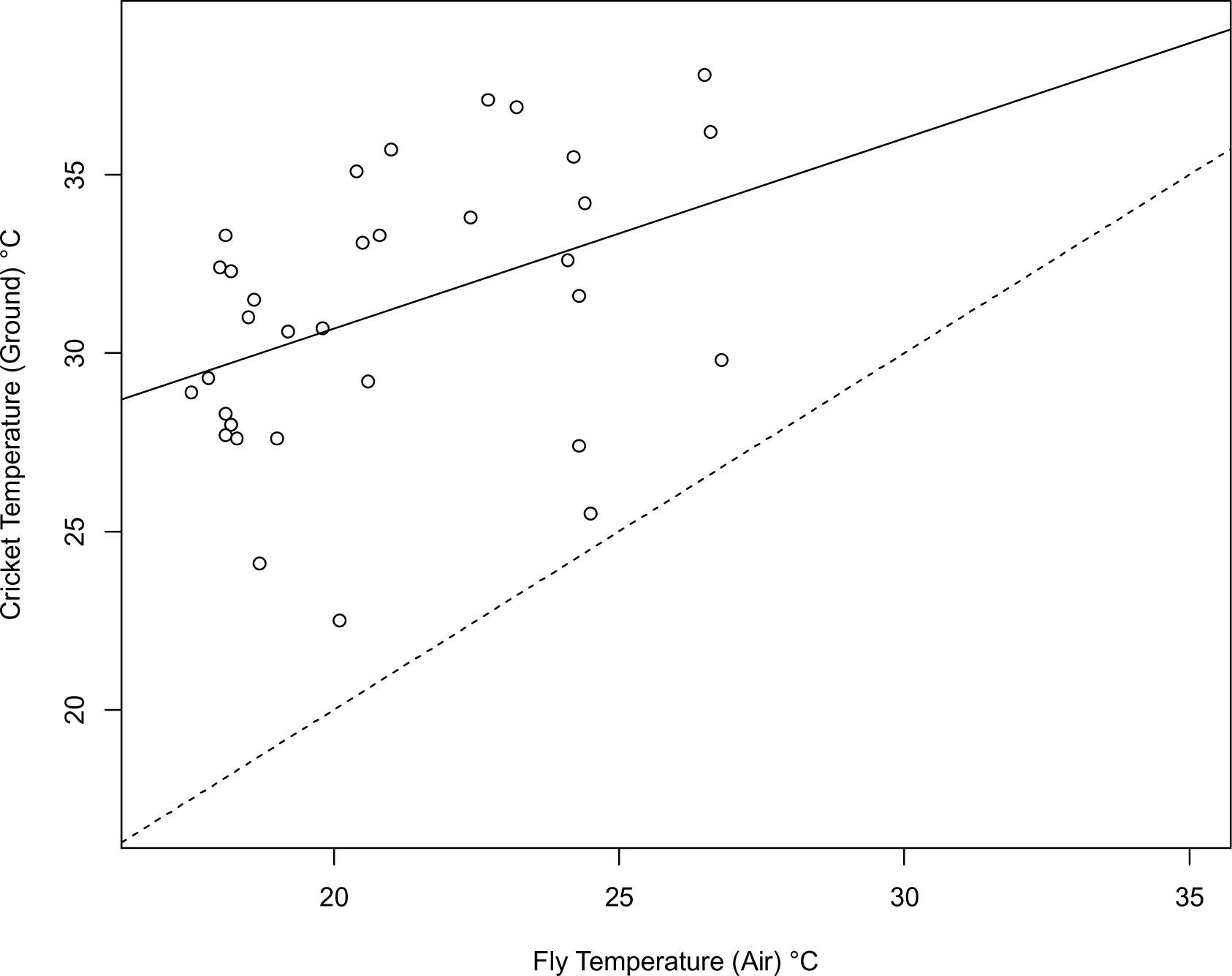
Correspondence between air temperature as experienced by female flies and ground temperature as experienced by male crickets. The solid line is the line of best fit to the data; it has a slope significantly greater than zero and significantly less than one (dashed line = equality).

## DISCUSSION

Here we show that temperature in the wild does indeed affect an eavesdropping parasitoid fly’s response to cricket mating signals, but that temperature induced variation in the signals themselves are not relevant to the fly, at least not within the ± 4 ℃ range tested here. The temperature effect on fly phonotaxis was to reduce the number of flies attracted to cricket song at temperatures below about 19 ℃ and above about 30 ℃. However, we note that we caught flies at all ambient temperatures from ∼15 to 35 ℃, so it is not simply the case that the ambient temperature range exceeded the physiological tolerances of *O. ochracea*. Our results are consistent with what we introduced as the Grandma scenario, and are inconsistent with fine tracking of temperature induced changes in cricket song pulse and chirp rates. On the one hand, this is surprising as previous work, also in the field with wild populations, has shown that *O. ochracea* shows a pretty high level of phonotactic specificity, demonstrating both geographic specialization favoring locally dominant host cricket songs (Gray et al., 2007), and even favoring certain song variants over others within a single local population (Gray and Cade, 1999; Sakaguchi and Gray, 2011; Wagner and Basolo, 2007). On the other hand, our data on the correspondence of air and ground temperatures suggest that precise song-temperature tracking by the fly is not especially practical, as air temperature had low predictive value for cricket ground temperature (R^2^ = 0.17), and ground temperature was not a set deviation from air temperature (slope not = 1).

Notably, we did not find a pattern of fly responses to cricket calls consistent with temperature coupling. This contrasts with the findings of Jirik et al. (2023) and Walker (1993) with a Florida population of the fly responding to variation in *G. rubens* pulse rates. There are a number of important differences between the California and Florida fly populations, and between our study design and that of Jirik et al. (2023) which may help interpret these differences. First, based on microsatellite DNA variation (Gray et al., 2019), it appears almost certain that the Florida population of flies has a longer co-evolutionary history with *G. rubens* than the California population of flies has with *G. lineaticeps*. In addition, the western USA populations of flies encounter and exploit a much wider range of host cricket species than do the Florida flies (Gray et al., 2019; Sakaguchi and Gray, 2011; Weissman and Gray, 2019). Furthermore, the western USA populations of flies, including the California population studied here, live in much drier climates than do the Florida flies. This is relevant in that the high humidity typical of Florida populations will tend to slow the rate of atmospheric cooling after sunset, such that the correspondence between air and ground temperatures in Florida is likely to be higher than in California. In the drier western USA populations, substantial microhabitat variation among potential cricket calling sites greatly reduces the air-ground correspondence, as does differential cooling of air and ground in the early evening hours when flies are most active (Beckers and Wagner, 2012; Cade et al., 1996). Taken together, evolutionary history and climactic differences make it plausible that Florida flies have evolved to recognize temperature specific pulse rates in *G. rubens* calls whereas California flies have not.

Alternatively, the California flies may respond to temperature induced changes in cricket song, but in ways we were unable to detect in the field given differences in experimental design. In their study, Jirik et al. (2023) used high resolution laboratory phonotaxis tests under controlled conditions of 21, 25 or 30 ℃. Their data clearly show maximal fly tracking of cricket pulse rates which match those expected for ambient, consistent with physiological temperature coupling. The experimental protocol they used is remarkably precise, allowing measurement of separate contributors to fly phonotaxis: walking distance and steering velocity. By comparison, our measures are extremely crude: the number of flies caught at the trap. The importance of our results is that they add the ecological context affecting natural selection in the wild: slight differences in walking distance and steering velocity observed in the laboratory may nonetheless all result in equivalent risk of parasitism in the field (indeed, we can not rule out that the flies we caught in the field differed slightly in their approach speed and/or efficiency dependent upon playback deviation from ambient). Given that temperature can vary drastically at small scales relevant to invertebrates (Jocson et al., 2019; Pincebourde et al., 2007; Pincebourde and Woods, 2012), it is likely that signalers and receivers/eavesdroppers regularly experience different thermal conditions. Overall, our data support a conclusion similar to that of Ritchie et al. (2001): physiological temperature coupling may be prevalent, but may not be highly relevant in the wild. This is especially likely to be the case for interspecific eavesdroppers as compared to conspecific males and females, and may vary among populations due to differences in ecology and/or evolutionary history.

## Contributor Roles

Conceptualization SMR, KDF-F, DAG; Data curation DAG; Formal Analysis DAG; Funding acquisition SMR; Investigation SMR, DAG; Methodology SMR, KDF-F, DAG; Visualization DAG; Writing – original draft SMR, DAG; Writing – review and editing SMR, KDF-F, DAG.

## Acknowledgments

Janelle Talavera, Stephanie Menjivar and Diego Gomez-Morales assisted with fieldwork. Norman Lee provided helpful feedback on an earlier draft of this article. Funding was provided by a Thesis Support grant from the CSUN Office of Graduate Studies to SMR.

## Data Availability

Analyses reported in this article can be reproduced using the data provided by Author (202x). [DRYAD upon article acceptance].

## Notes

### Competing Interest Statement

The authors have declared no competing interest.

